# Spatial transcriptomic analysis reveals coordinated gene expression in ovarian clear cell carcinoma and adjacent endometriosis in UK and Japanese patients

**DOI:** 10.64898/2026.05.29.728698

**Authors:** Takafumi Kuroda, Gaia Giannone, Darren P. Ennis, Hasan B. Mirza, Daniel Marks, Louis Flood, Mary Sisley, Ryan Griffin, Saral Desai, Jacqueline McDermott, Nathalie Lambie, Nei Fukasawa, Takako Kiyokawa, Masayuki Shimoda, Misato Saito, Takuhiro Koba, Ryosuke Saito, Ayako Kawabata, Masataka Takenaka, Giorgio Valabrega, Nik Matthews, Laura A Tookman, Nozomu Yanaihara, Aikou Okamoto, Iain A. McNeish

## Abstract

**Purpose:** Ovarian clear cell carcinoma (OCCC) is strongly associated with endometriosis and shows geographic variation in incidence. We investigated whether OCCC and adjacent endometriosis exhibit distinct transcriptional states and whether these patterns differ between United Kingdom (UK) and Japanese cohorts.

**Experimental Design:** We performed whole-transcriptome spatial profiling on specimens from 16 OCCC cases (8 UK, 8 Japan) in which tumor and endometriosis were both present. Gene expression was analyzed in tumor, endometriosis and stroma. ARID1A status was assessed by immunohistochemistry.

**Results:** Median age was 59 years (range 26–82). 13/16 cases (81.3%) had early-stage disease. Tissue compartment rather than cohort of origin was the dominant source of variation across endometriosis and tumor regions. Endometriosis was enriched for inflammatory and immune-related pathways compared to tumor, whilst there was greater representation of chromatin and protein–DNA complex assembly pathways in tumor regions. These patterns were conserved across both cohorts and after stratification by ARID1A status. Mesenchymal-associated gene expression scores also significantly differed across stroma, endometriosis and tumor with clear compartmental separation. Cell type deconvolution analyses showed clear compositional differences between stromal and epithelial disease compartments.

**Conclusions:** OCCC and coexisting endometriosis are transcriptionally distinct, with the dominant contrast being compartmental rather than geographic. ARID1A alone is unlikely to account for the principal spatial transcriptional states identified here. Further analyses will be required to ascertain whether these differences reflect genuine biological differences between OCCC and coexisting endometriosis or represent different stages of endometriosis-associated tumorigenesis.

**Translational Relevance:** Ovarian clear cell carcinoma often arises in association with endometriosis, yet the biological transition between these lesions remains poorly understood. Using spatial transcriptomics in matched tumor and adjacent endometriosis from Japanese and UK cohorts, we showed that endometriosis is characterized by inflammatory and antigen-presentation features, whereas tumor regions showed chromatin-organization and oncogenic transcriptional states. These patterns were largely maintained irrespective of ARID1A status and geographic background. In addition, spatial deconvolution suggested differences in local immune composition, with tumor regions showing relatively greater neutrophil- and T cell-associated signals. Together, our data suggest that OCCC and coexisting endometriosis share a spatially linked tissue context, but that tumor regions have distinct transcriptional profile and microenvironment that may be involved in the malignant transformation and inform interpretation of molecular classification in endometriosis-associated OCCC.

## Introduction

Ovarian clear cell carcinoma (OCCC) is a rare disease, representing approximately 10% of all ovarian cancers (1,2). Its incidence is significantly higher in Southeast Asian populations (10–30% compared with 5–12% in Western populations), although the underlying reasons for this difference remain unclear (3,4). OCCC is typically diagnosed at an early stage, with good prognosis; survival rates are around 88% for stage I and 70% for stage II disease (3). However, outcomes are extremely poor for advanced stage or recurrent OCCC, largely due to inherent chemoresistance and aggressive biological behavior (3). OCCC has distinct clinical features, molecular profile and disease course compared to the more common high-grade serous carcinoma, and in several respects more closely resembles clear cell renal carcinoma. This distinction has led to recognition of OCCC as a discrete disease entity requiring histotype-specific investigation and management (5,6).

Evidence suggests that endometriosis is not only a significant risk factor but may also act as a precursor lesion (7,8). This is supported by pathological series identifying endometriosis in approximately half of OCCC cases and by population-based data showing a three-fold age-adjusted increase in the likelihood of developing OCCC in women with histologically proven endometriosis (9). Endometriosis is a highly inflammatory condition, marked by activation of IL-6– related signaling, altered macrophage and NK-cell functional states and concurrent impairment of tumor-suppressor mechanisms (10,11). Mutations in *KRAS, PIK3CA* and *ARID1A*— the latter altered in approximately half of OCCC cases — have been identified across normal epithelium, endometriotic tissue and OCCC, further supporting a stepwise evolutionary process (12-14), and the mutations observed in OCCC and adjacent endometriosis overlap closely, with none unique to the carcinoma (15). However, although loss of ARIDA1 function is the most frequent recurrent alteration in OCCC, it does not appear to have a prognostic role (16-18).

Technological advances, particularly in single-cell transcriptomics, have provided insight into the molecular and cellular landscape of endometriosis, reinforcing the concept of a continuum from the common benign endometriosis and the rare and aggressive OCCC (19,20). However, no study to date has directly compared spatially mapped, lesion-adjacent endometriosis and matched OCCC from the same patients using approaches that combine spatial transcriptomics with precise topographical mapping of the tissue architecture.

For this reason, we conducted a study comparing OCCC and associated endometriosis in two clinically comparable patient cohorts from Japan and the UK, leveraging an international dataset to explore conserved and population-specific features underlying OCCC pathogenesis.

## Methods

### Patients and samples

Patients diagnosed with ovarian clear cell carcinoma (OCCC) between 2017 and 2023 at Imperial College Healthcare NHS Trust, London (UK cohort) and The Jikei University Hospital, Toyko (Japanese cohort - JP) were included in this study. Samples from the UK cohort were used under the authority of the Imperial College Healthcare Tissue Bank (HTA license 12275, ethics reference 22/WA/0214, Project number R21057). Samples from the JP cohort were provided by The Jikei University School of Medicine and the study protocol was approved by the local Ethics Review Committee [37-052 (12689)]. A data transfer agreement (DTA) was established between the institutions for data sharing.

### Pathological assessment

Cases were deemed eligible when both tumor and endometriosis lesions were present within the same formalin-fixed paraffin-embedded (FFPE) tissue block. All cases underwent pathological re-review by expert gynecologic pathologists and the eight most recent cases each from the UK and JP were selected. Thereafter, a further a 4μm section underwent H&E staining and was reviewed immediately before cutting the section used for spatial transcriptomic analysis to guarantee the quality and consistency of the sample.

### ARID1A immunohistochemistry

ARID1A status was assessed by immunohistochemistry using institution-specific validated protocols. In the UK cohort, staining was performed using ARID1A antibody clone EPR13501 (ab182560, Abcam) at a dilution of 1:1000 according to the published protocol (21). In the JP cohort, staining was performed using a polyclonal anti-ARID1A antibody (NBP1-88932; Novus Biologicals) at a dilution of 1:200, as previously described (22). ARID1A loss was defined as complete absence of nuclear staining in tumor cells, with retained nuclear staining in stromal cells serving as an internal control.

### Sample and library preparation

Unstained FFPE sections were prepared for GeoMx Digital Spatial Profiler (DSP) analysis using the Human Whole Transcriptome Atlas (WTA) RNA panel. Slide preparation was performed using a semi-automated workflow with the BOND RX system (JP cohort) and a manual workflow (UK cohort) according to the manufacturer’s recommended protocol. After deparaffinization and heat-induced target retrieval followed by protease treatment, WTA probes were hybridized to the tissue sections.

### ROI selection

Regions of interest (ROIs) were defined jointly by a researcher and a pathologist after overlapping the corresponding H&E and immunofluorescence images and annotated based on morphological features as well as staining patterns of DNA (DAPI), pan-cytokeratin (PanCK) and CD45. ROIs were classified as stroma, endometriosis, tumor, tumor–stroma interface, endometriosis-passage (EP) and tumor-passage (TP). EP and TP were defined as transitional regions identified under pathologist supervision; although morphologically classified as endometriosis or tumor, respectively, they were located at the boundary with adjacent tissues. Thus, these regions were not treated as independent lesions, but rather as transitions toward the neighboring tissue.

### Sequencing and initial data processing

Photo-cleaved oligonucleotide tags released from each ROI were collected into dedicated collection plates and stored until downstream library preparation. For NGS readout, collected oligonucleotide tags were amplified by PCR using the GeoMx Seq Code system and the resulting libraries were pooled and purified. Library concentration and fragment quality were assessed using the Qubit dsDNA HS Assay and Agilent TapeStation.

Libraries were prepared using the GeoMx DSP NGS Readout protocol with GeoMx Seq Code primer plates and processed on an Illumina NextSeq 2000 platform. Raw sequencing data were generated in FASTQ format and converted to digital count conversion (DCC) files using the GeoMx NGS Pipeline. The resulting DCC files were uploaded to the GeoMx system and then exported for downstream quality control and expression analyses were performed using R software ver.4.4.2.

### Differential expression analysis

Before integrated analysis of the UK and JP cohorts was performed, batch correction was performed using ComBat (23), as detailed in Supplementary Methods. Differential expression analysis was performed using limma on the ComBat-corrected expression matrix. Linear models were fitted using lmFit and moderated statistics were obtained using contrasts.fit and eBayes. Each ROI was treated as an independent observation, consistent with prior ROI-level spatial transcriptomic analyses (24). Differentially expressed genes (DEGs) were defined as those meeting both false discovery rate (FDR) < 0.05 and an absolute log2 fold change of ≥1.

The primary analysis compared tumor ROIs with endometriosis ROIs. In addition, to evaluate cohort-related expression tendencies within each compartment, UK and JP were compared separately within tumor ROIs and within endometriosis ROIs. ARID1A-stratified differential expression analyses were also performed.

### Pathway analysis and fixed pathway comparison

Pathway enrichment analysis of DEGs was performed using clusterProfiler. Over-representation analysis was performed for both Gene Ontology Biological Process (GO:BP) and KEGG. For the primary tumor versus endometriosis analysis, enrichment analyses were performed independently for the combined dataset (UK+JP), the UK cohort alone and the JP cohort alone. For fixed pathway comparison, among pathways significantly enriched in the combined analysis, the top 10 terms ranked by FDR were selected separately for tumor-upregulated and endometriosis-upregulated analyses and these were defined as the fixed term list. For both GO:BP and KEGG, this combined-derived fixed term list was used as a common reference axis and enrichment results from the UK and JP cohorts were compared across the same set of terms.

### Clinical and pathological variables

For each case, age, stage, tumor ARID1A status, total number of ROIs and the numbers of ROIs corresponding to tumor and endometriosis were recorded.

### Statistical analysis

All analyses and plot generation were performed using R version 4.4.2. Continuous variables were compared using nonparametric tests and categorical variables using Fisher’s exact test. To account for multiple testing, *P* values were adjusted using the Benjamini–Hochberg method and pathways with FDR <0.05 were considered significantly enriched.

## Results

### Clinical features

Patients with a diagnosis of OCCC between 2017 and 2023, treated at Imperial College Healthcare NHS Trust, London, UK and the Jikei University Hospital, Tokyo, Japan (JP) were reviewed. A representative H&E section was reviewed by two expert pathologists. All samples included had OCCC and adjacent endometriosis in the same FFPE block ensuring that both tumor and endometriosis were represented in the same section. Sixteen OCCC formalin-fixed paraffin-embedded (FFPE) blocks were selected, eight cases from each cohort (hereafter UK and JP cohorts – Figure 1A). The main clinical features are summarized in Table 1. Median age at diagnosis was 59 years (range 26–82), and most cases were FIGO 2014 stage I or II (81.3% (13/16)), with only three advanced-stage cases (III/IV, 18.7%). Stage distribution (I/II vs III/IV) was similar between cohorts (Table 1). ARID1A loss by IHC was observed in 37.5% (6/16) tumors overall (JP cohort 5/8; UK cohort 1/8 with no statistical difference between cohorts) (Table 1).

**Table 1.** Clinicopathologic characteristics and spatial sampling. Clinicopathologic characteristics and ROI distribution in the combined, UK, and JP cohorts. Sixteen cases were included (UK, n = 8; JP, n = 8). Median age (range), stage (I/II or III/IV), ARID1A status in tumor tissue (retained or loss), total ROI number, ROI counts for tumor and endometriosis and median (range) ROIs per case are shown. *P* values were calculated using the Mann–Whitney test (a) or Fisher’s exact test (b), as indicated.

| Characteristics and spatial sampling | Total | Cohort |  | *P*-value |
| --- | --- | --- | --- | --- |
|  |  | UK | JP |  |
| Number of cases | 16 | 8 | 8 |  |
| Age<br>Median (range) | 59 (26-82) | 64 (26-82) | 57 (54-75) | 0.91<sup>a</sup> |
| Stage<br>I/II<br>III/IV | 13<br>3 | 7<br>1 | 6<br>2 | 1.0<sup>b</sup> |
| ARID1A expression in tumor<br>Retained<br>Lost | 10<br>6 | 7<br>1 | 3<br>5 | 0.12<sup>b</sup> |
| ROI numbers<br>Tumor - total<br>Endometriosis - total<br>Median (range) per case | 280<br>62<br>60<br>17 (13-22) | 136<br>30<br>27<br>16 (13-22) | 144<br>32<br>33<br>18 )17-19) |  |

**Figure 1.**
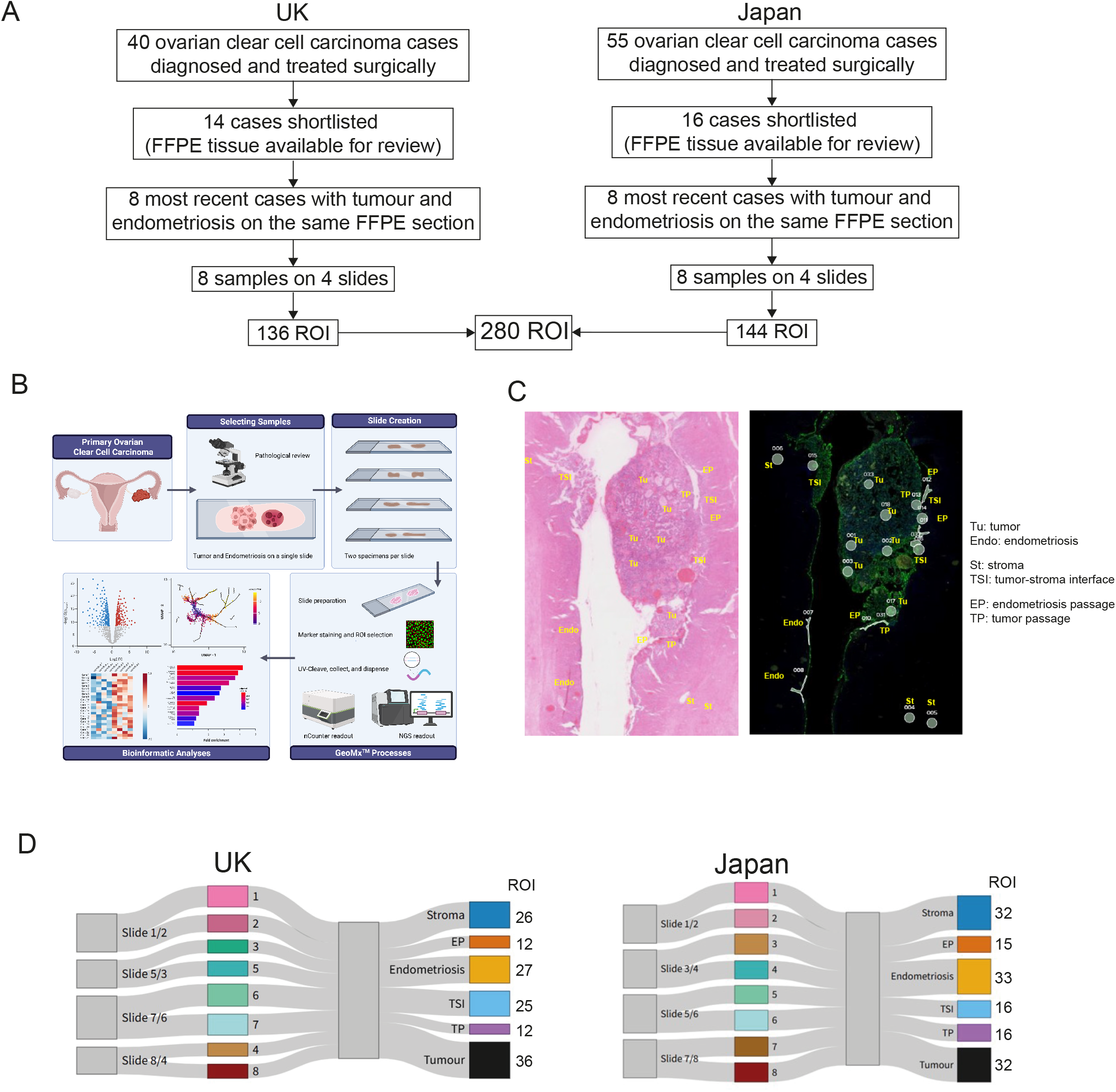
Patients, spatial profiling and data quality. **A**. Sample selection workflow for the UK and JP cohorts. Eight cases were selected in each cohort. A total of 136 ROIs were profiled in the UK cohort and 144 in the JP cohort. **B**. Study design and GeoMx workflow. **C**. ROI selection and marker-based visualization of tumor and endometriosis regions. Tumor (Tu), endometriosis (Endo), stroma (St), tumor–stroma interface (Tu–St inter), endometriosis-passage (E–P), and tumor-passage (T–P) regions are shown. The corresponding immunofluorescence image shows DNA (blue), PanCK (green), and CD45 (red). Circular overlays indicate selected ROIs. **D**. Sankey-style diagrams showing the distribution of profiled cases and ROI categories in the UK (left) and Japan (right) cohorts.

### Study design and workflow

The overall study design and GeoMx workflow are shown in Figure 1B. Regions of interest (ROIs) were defined on matched sections with reference to the corresponding H&E-stained slide and GeoMx immunofluorescence images using morphology markers (PanCK, CD45 and DAPI). ROI selection was performed by a researcher and an expert pathologist on the basis of morphology and marker staining (Figure 1C). ROIs were sampled across tumor (Tu), endometriosis (Endo), stroma (St), tumor–stroma interface and passage regions, yielding a total of 280 ROIs (UK, 136; JP, 144; Figure 1A and Table 1) as described in the Methods. ROI distribution by cohort and compartment is summarized in Figure 1D and Table 1.

Quality control (QC) and detection metrics were comparable between cohorts. After QC filtering, analyzable ROIs were obtained for nearly all selected regions (99.6% 277/280), with a negligible number of QC warnings (Table S1). A similar number of endogenous genes were detected above the limit of quantification in both cohorts (UK: 13,595/18,677; JP: 15,612/18,677). Gene detection above the limit of quantification decreased with increasing segmentation stringency in both cohorts (Table S1).

### No cohort effect when comparing the UK and Japanese cases

A critical first question was whether there were significant differences between cases from UK and Japan. Separate analyses of the cohorts showed that ROIs clustered largely according to ROI compartment annotations, with stromal ROIs forming a particularly distinct cluster (Figure S2). When the two cohorts were initially integrated, cohort-specific clustering became evident (Figure 2A). However, after batch correction, cohort mixing increased (Figure 2B), accompanied by reduced cohort association of housekeeping-gene expression (Figure S3) and improved mixing in stromal ROI PCA analyses (Figure 2D).

**Figure 2.**
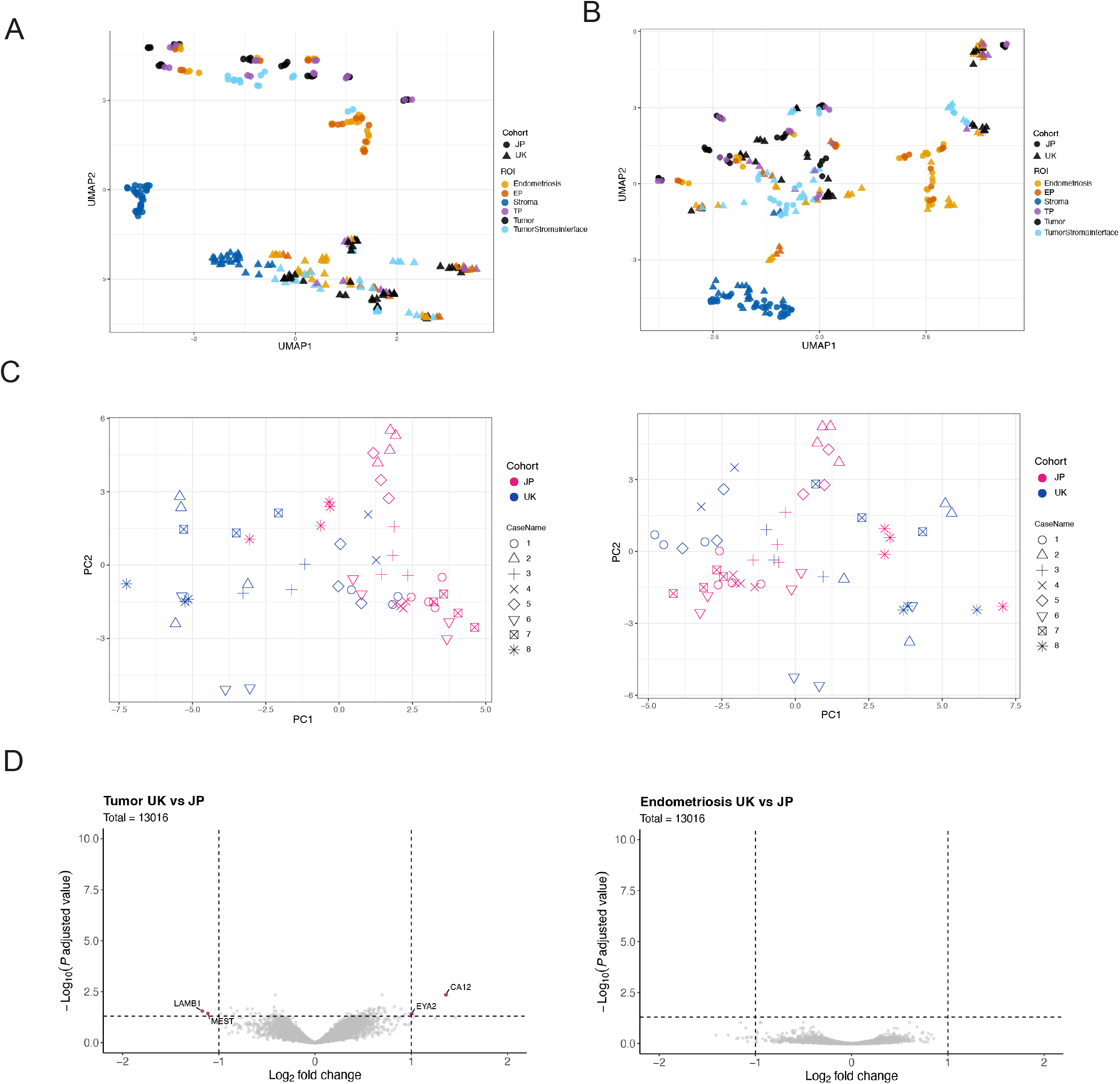
Cohort effects after batch correction. **A, B**. Unsupervised clustering of ROIs visualized by UMAP before (**A**) and after (**B**) batch correction. Each point represents one ROI and is colored by tissue category. Cohort origin is indicated by symbol shape. **C**. PCA of stromal ROIs before (left) and after (right) correction. Points are colored by cohort and shaped by case. **D**. Volcano plots showing differential gene expression between the UK and Japan (JP) cohorts after ComBat correction. The left panel shows tumor ROI and the right panel shows endometriosis ROIs. The x-axis represents log2 fold change (JP vs UK) and the y-axis represents −log10 adjusted P value. Dashed lines indicate fold-change and adjusted *P* value thresholds.

In the corrected dataset, differential expression between the UK and JP cohorts was very limited in both tumor and endometriosis ROIs, with few genes meeting significance thresholds (Figure 2D). Consistent with this, pathway enrichment analyses did not identify any consistent cohort-specific pattern. Taken together, these results suggest that cohort-of-origin was not a major source of variation after correction and that gene expression in UK and Japanese OCCC is very similar. As a result, subsequent analyses were performed on the combined batch-corrected dataset.

### Tumor and adjacent endometriosis have different transcriptional profile and heterogeneity related programs

We first explored transcriptional differences between tumor and endometriosis, hypothesizing that, although genomically similar, they have differences in the gene expression profile. Differential expression analyses identified differentially expressed genes (DEGs) between tumor and endometriosis (Figure 3A). Using thresholds of FDR <0.05 and |log2 fold change| ≥1, 144 genes were upregulated in endometriosis and 58 genes were upregulated in tumor (Figure 3B). DEGs in endometriosis included inflammatory and immune-associated transcripts such as *IL1R1* and *C4B*, as well as antigen presentation–related transcripts (Figure 3A and Table S2). In tumor areas, DEGs included upregulation of histone/chromatin-associated transcripts such as *H2BC17, H2AC17, H3C13* and *H3C15* (Figure 3A). Among tumor-upregulated genes, 24 were shared between the UK and JP cohorts (42 and 110 genes, respectively). Among endometriosis-upregulated genes, 74 were shared between the two cohorts (97 and 278 genes, respectively) (Figure 3B). Shared tumor-upregulated genes, including *H1-5, H3C10* and *H2BC17*, suggested a proliferative and chromatin-active tumor state. Shared endometriosis-upregulated genes, including *HLA-DRA, CD74, CXCL8, CCL2, PTGS2* and *C3*, suggested an inflammatory, immune-active state with a prominent MHC class II antigen-presentation component. The full list of shared genes is provided in Table S2.

**Figure 3.**
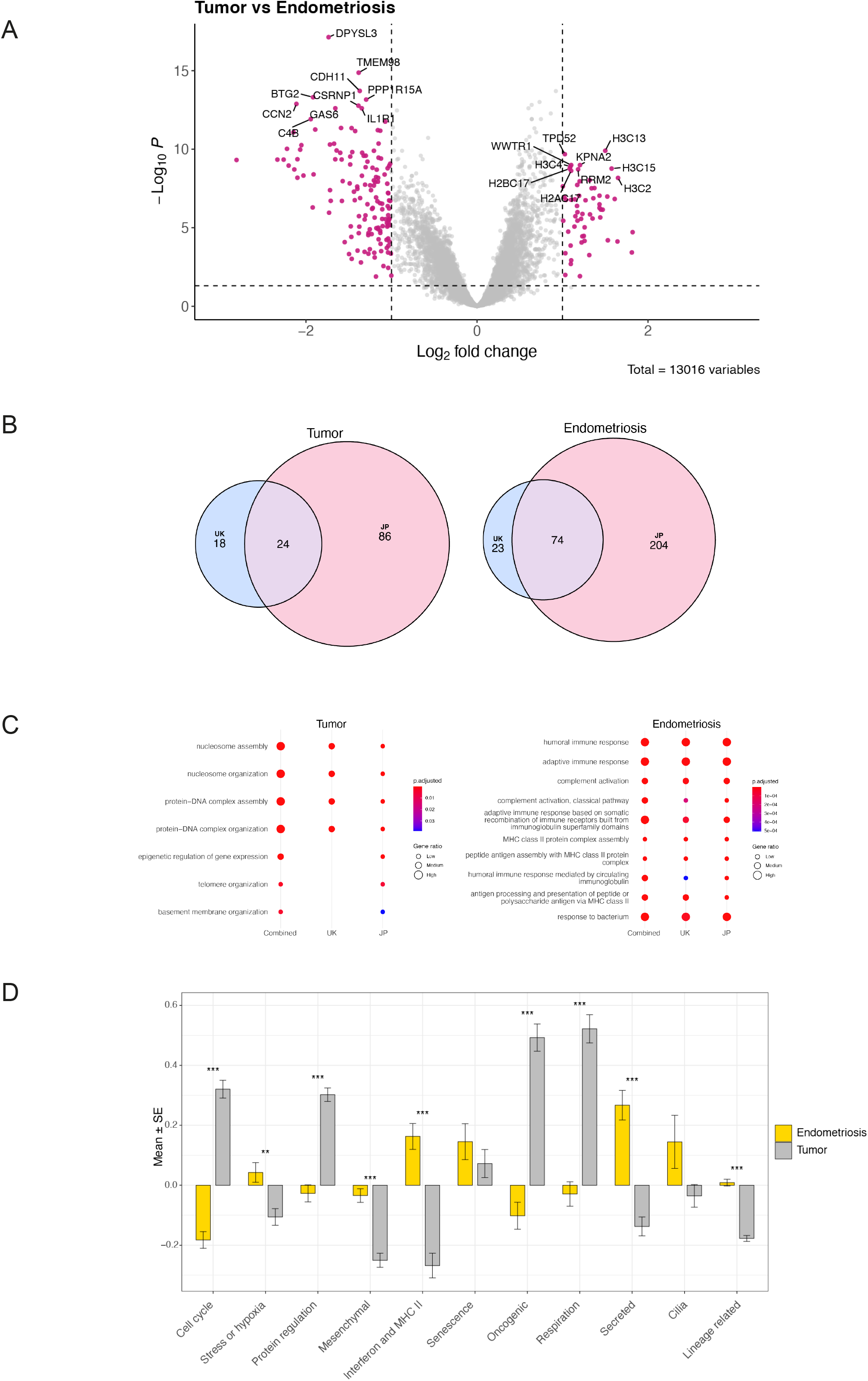
Transcriptional differences between tumor and endometriosis. **A**. Volcano plot showing differential gene expression between tumor and endometriosis ROIs after batch correction. The x-axis represents log2 fold change (tumor vs endometriosis) and the y-axis represents −log10 adjusted P value. Dashed lines indicate |log2 fold change| = 1 and FDR = 0.05. Genes meeting both thresholds are highlighted. **B**. Venn diagrams showing the overlap of differentially expressed genes between the UK and Japan cohorts. Panels show genes upregulated in tumor relative to endometriosis (left) and gene upregulated in endometriosis relative to tumor (right). Numbers indicate cohort-specific and shared DEGs. **C**. GO Biological Process enrichment for tumor-upregulated (left) and endometriosis-upregulated (right) gene sets across the combined, UK and Japan analyses. The same term list defined from the combined analysis is shown for each cohort-specific analysis to facilitate comparison. Dot size indicates gene ratio and color indicates adjusted *P* value. **D**. Category-level summary of meta-program scores in tumor and endometriosis ROIs after batch correction. Bars indicate mean ± SE. Category-wise comparisons were performed using two-sided Wilcoxon rank-sum tests with Benjamini–Hochberg correction. Significance is indicated by FDR-adjusted *P* values. *FDR < 0.05, ** FDR < 0.01, *** FDR < 0.001.

Across cohort-stratified analyses, Gene Ontology Biological Process (GO:BP) enrichment patterns were comparable among the combined, UK and JP cohorts (Figure 3C), with similar results using KEGG (Figure S4). Tumor-upregulated pathways were mainly related to chromatin- and genome-organization–related processes, including nucleosome assembly/organization and protein–DNA complex assembly/organization (Figure 3C; Figure S3). Conversely, endometriosis-upregulated pathways were mainly linked to immune and inflammatory programs, including complement activation and immune activation–related terms (Figure 3C; Figure S3). Together, these data indicate that tumor and endometriosis ROIs have very distinct transcriptional programs, with endometriosis being characterized by a proinflammatory gene expression profile, while tumor areas upregulate mainly genes involved in chromatin accessibility and epigenetic remodeling. Similar trends were observed in all analyses when the UK and JP cohorts were analysed independently before batch correction (Figure S5, Table S2-4).

In view of the potential of a gradual change in gene expression from endometriosis to tumor, we used the spatial information from GeoMx to define and mark areas of endometriosis at the interface with tumor (which we termed endometriosis-passage) and the corresponding cancer areas (which we termed tumor-passage). When comparing endometriosis and endometriosis– passage, we did not detect any difference in DEGs; equally, the results were very similar when comparing tumor and tumor–passage (Figure S6).

To validate our findings in the context of the broader tissue landscape, compartment-level comparisons against stromal ROIs were also performed and showed clear transcriptional differences for both endometriosis and tumor relative to stromal ROIs (Figure S7). To resolve the broad differences between stroma, endometriosis and tumor ROIs further, we used the previously reported 41 pan-cancer meta-program (MP) gene sets, grouped into 11 hallmarks of intratumor heterogeneity (25), as predefined signatures for ROI-level scoring of our data. This provided a useful framework for exploring tumor–endometriosis crosstalk in addition to conventional differential expression analysis.

The results again demonstrate that there are distinct expression programs across stroma, endometriosis and tumor (Figure S8), with tumor showing enrichment for oncogenic and cell cycle– related MPs, whereas endometriosis showed higher immune-related MPs, including interferon and MHC II (Figure 3D). Multiple MPs remained significant after correction for multiple testing (Figure 3D).

### The endometriosis-to-tumor transcriptional shift was largely preserved across ARID1A groups

ARID1A status was assessed by immunohistochemistry, with loss defined as complete absence of nuclear staining in tumor cells with retained stromal internal controls (Figure 4A; Table 1). *ARID1A* transcript abundance did not differ significantly between ARID1A-loss and ARID1A-retained tumor ROIs, nor between UK and Japan samples (Figure S9), in keeping with previous reports (26).

**Figure 4.**
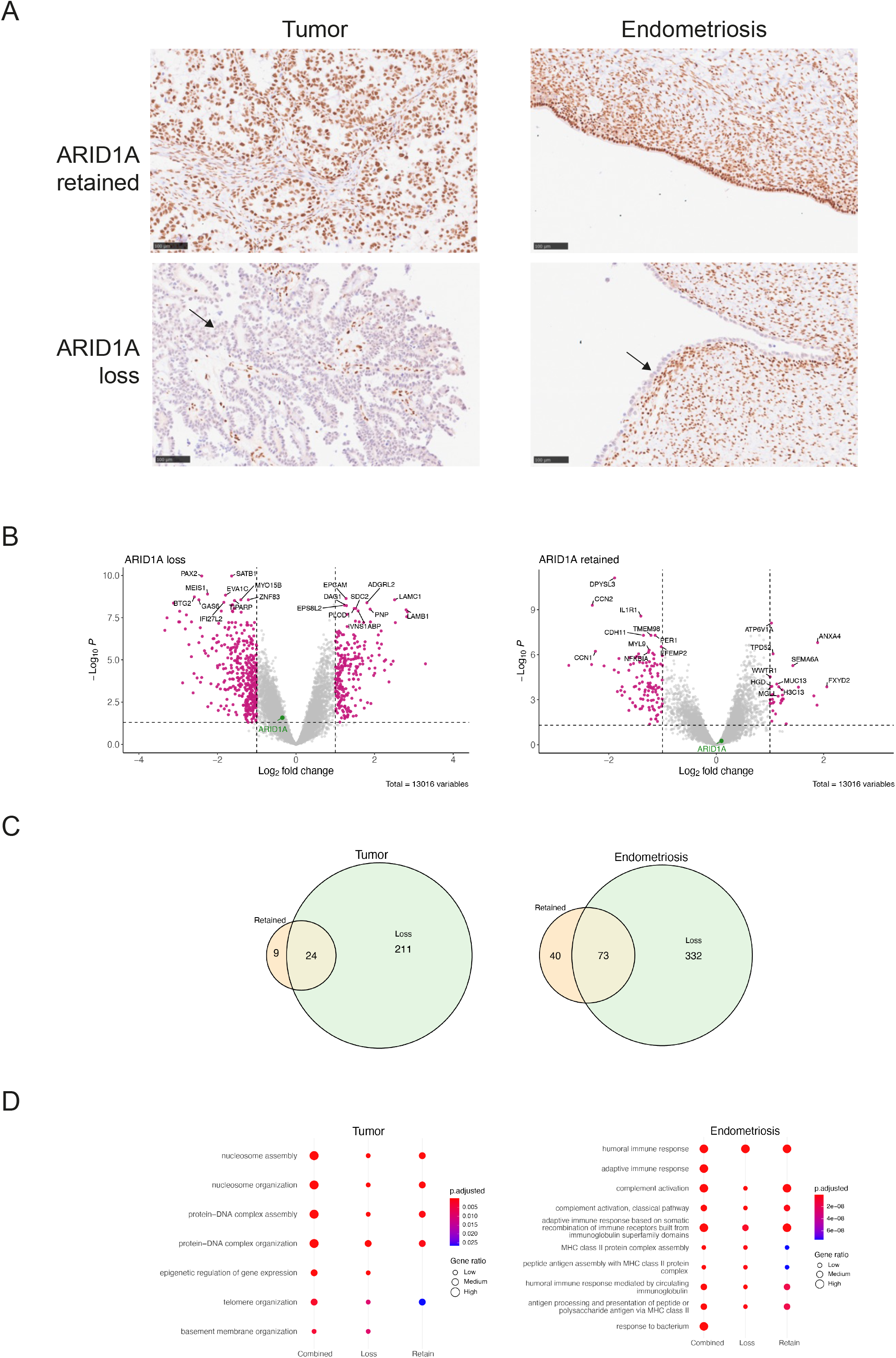
ARID1A stratification and tumor–endometriosis differences. **A**. Representative immunohistochemical staining for ARID1A in tumor (left) and endometriosis (right) ROIs with retained (upper panels) and absent (lower panels). ARID1A loss was defined as complete absence of nuclear staining in epithelial cells (arrows) with retained stromal internal controls. Scale bar - 100 μm. **B**. Tumor–endometriosis differential expression repeated within ARID1A-retained (left) and ARID1A-loss (right) samples. Volcano plots show the distribution of differentially expressed genes in each stratum. The x-axis represents log2 fold change (tumor vs endometriosis) and the y-axis represents −log10 adjusted P value. Dashed lines indicate |log2 fold change| = 1 and FDR = 0.05. **C**. Overlap of tumor-upregulated and endometriosis-upregulated gene sets between ARID1A-retained (left) and ARID1A-loss (right) samples. Shared genes are indicated in the intersection. The left panel shows tumor-upregulated genes and the right panel shows endometriosis-upregulated genes. **D**. GO Biological Process enrichment for tumor-upregulated and endometriosis-upregulated gene sets across the combined and ARID1A stratified analyses. The same term list defined from the combined analysis is shown for each cohort-specific analysis to facilitate comparison. Dot size indicates gene ratio and color indicates adjusted *P* value.

We next performed tumor-versus-endometriosis comparisons stratified by ARID1A status (Figure 4B,C). Of the tumor-upregulated genes, 24 were shared between the two groups (out of 33 genes in the ARID1A-retained group and 235 in the ARID1A-loss group). Of the endometriosis-upregulated genes, 73 were shared (out of 113 genes in the ARID1A-retained group and 405 in the ARID1A-loss group) (Figure 4B and 4C; Table S6). Shared tumor-upregulated genes, including *ANXA4, WWTR1, H3C10* and *H3C13*, were consistent with a proliferative and chromatin-active tumor state, whereas shared endometriosis-upregulated genes, including *HLA-DRA, CD74, CXCL8, CCL2* and *C3*, indicated an inflammatory, immune-active state with a prominent MHC class II antigen-presentation component. The full list of shared genes is provided in Table S6.

We then analyzed upregulated pathways between these two regions stratifying for ARID1A status. In both groups, tumor-upregulated pathways were enriched for chromatin-related processes, including nucleosome assembly/organization and protein–DNA complex assembly/organization (Figure 4D, Figure S11, Table S6). Similarly, pathways related to immune activation were upregulated in endometriosis (Figure 4D, Figure S10, Table S6). Both groups demonstrated concordant directional changes in pathway activity irrespective of ARID1A status, with only modest differences in the degree of alteration.

### There are microenvironmental differences across compartments with a higher proportion of neutrophils and T lymphocytes in tumor

We next evaluated the published EpiCC/MesCC framework, a bulk transcriptomic OCCC classification associated with prognosis and potential therapeutic stratification (27). Because this framework was originally defined using bulk expression data, we first confirmed differences in scores across the whole cohort (Figure S11). We then applied it to ROI-level profiles as a continuous MesCC score rather than for discrete case reassignment. MesCC scores differed significantly across tissue compartments, with the highest values in stroma, intermediate values in endometriosis and the lowest values in tumor ROIs (Figure 5A). These findings suggest that the bulk-derived MesCC signal may largely reflect gene expression in non-tumor compartments, especially the proportion of stromal cells within samples.

**Fig. 5.**
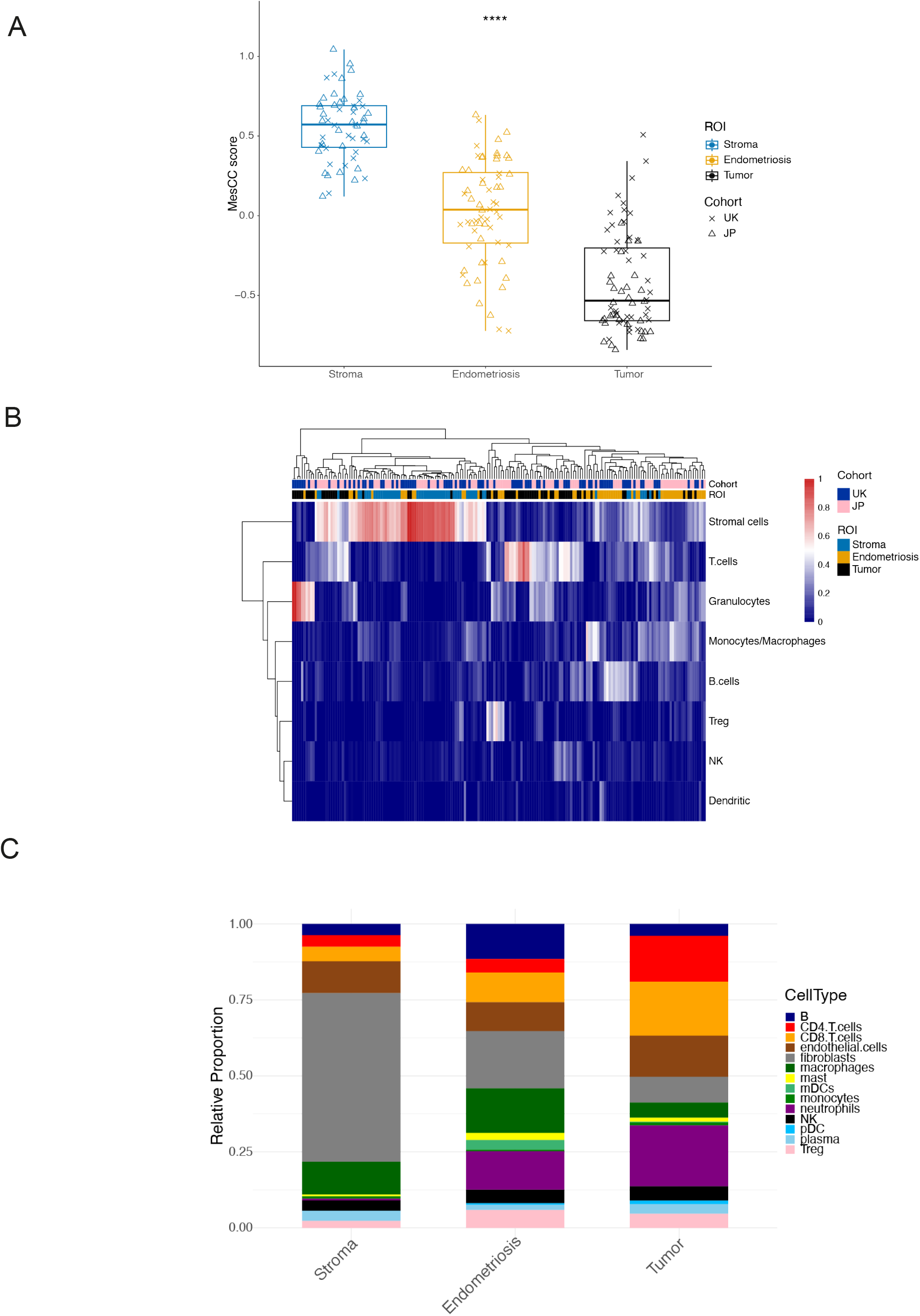
Microenvironmental differences across compartments. **A**. MesCC scores in stroma, endometriosis, and tumor ROIs after batch correction. Points represent individual ROIs. Symbols indicate cohort. Group differences were assessed using the Kruskal–Wallis test. ****; *P*<0.0001. **B**. Heatmap showing SpatialDecon-estimated non-tumor cell-type proportions across stroma, endometriosis, and tumor ROIs. Columns represent individual ROIs and rows represent merged cell types. **C**. Relative non-tumor cell-type composition in stroma, endometriosis and tumor ROIs. For each tissue type, cell-type values were summed across all ROIs and expressed as the fraction of the total signal across all cell types.

This prompted us to define the spatial distribution of cell types across stroma, tumor and endometriosis using a validated pipeline, SpatialDecon, which was developed specifically for GeoMx (28). We adapted the provided cell profile matrix and incorporated the gene expression profiles of representative tumor ROIs to account for tumor cells and then estimated the proportions of non-tumor cell populations using the cell-type definitions shown in Table S8. The resulting heatmap showed no obvious clustering by UK and JP cohort (Figure 5B).

Estimated cell proportions differed across compartments (Figure 5C, Figure S13). Stromal ROIs were characterized by high fibroblast-associated signals, whereas endometriosis and tumor ROIs showed higher proportions of immune-related cell types (Figure 5B and 5C). Tumor and endometriosis ROIs also appeared to differ in immune composition, with tumor ROIs showing relatively higher neutrophil- and T cell-associated signals, including CD4 and CD8 T-cell populations, whereas endometriosis ROIs showed relatively higher macrophage-, B-cell- and myeloid dendritic cell-associated signals, in keeping with the broader enrichment of antigen-presentation-related immune features in this compartment (Figure 5C, Figure S12 and Table S8). Similar biological patterns were observed in both the JP and UK cohorts (Figure S13).

## Discussion

In this study, we used spatial transcriptomic analysis to compare ovarian clear cell carcinoma (OCCC) and coexisting endometriosis in 16 cases in which both components were present on the same FFPE section. The principal aim of this study was to assess the major transcriptional differences between tumor and endometriosis and whether any such patterns differed between UK and Japanese patients. A secondary aim was to explore the possible contribution of ARID1A status to these spatial transcriptional states.

We showed that there was a negligible difference in the gene expression profile between the two cohorts and variation reflected tissue compartment, namely tumor versus endometriosis, rather than cohort of origin. This is an important observation because, to our knowledge, there have been no previous spatial transcriptomic studies directly comparing OCCC and adjacent endometriosis across clinically comparable cohorts from different geographic settings. This also has implications for the conduct of clinical trials in OCCC, facilitating recruitment across continents and interpretation of multi-national datasets.

The most consistent biological difference in this study was therefore the contrast between tumor and adjacent endometriosis. Genes upregulated in endometriosis were enriched for inflammatory and immune-related programs, including complement activation, antigen presentation and MHC class II-related transcripts. In contrast, genes upregulated in tumor were enriched for programs related to nucleosome assembly, chromatin organization and protein–DNA complex organization. This pattern was observed not only in the integrated analysis but was also broadly reproduced in cohort-specific analyses. Taken together, these findings support the view that OCCC and coexisting endometriosis are biologically related but represent distinct transcriptional evolutions within the same process.

Meta-program analysis further supported this compartment-centered interpretation (25). Tumor regions showed stronger enrichment in oncogenic and cell cycle-related programs, whereas endometriosis showed stronger enrichment of immune-related programs centered on interferon signaling and MHC class II. Although GeoMx does not provide single-cell resolution, the concordance between differential expression analysis and meta-program analysis strengthens the interpretation that the tumor–endometriosis difference in this dataset reflects coordinated biological programs rather than isolated marker genes alone.

*ARID1A* mutation is an important molecular alteration in OCCC and has also been implicated in endometriosis-associated tumorigenesis (12,13,19). In our cohort, the transcriptional contrast between tumor and endometriosis was broadly maintained after stratification by ARID1A status. Thus, in the present dataset, ARID1A status alone did not appear to account for the major changes and transcriptional evolution that were identified when comparing tumor and endometriosis, supporting the view that *ARID1A* is not a simple driver of the established OCCC phenotype. Rather, it is one factor within a broader biological context that may include chromatin remodeling, inflammatory signaling and local tissue environment.

Given the very limited availability of gene-expression classifiers specifically developed for OCCC, we evaluated the MesCC score as an OCCC-relevant bulk-derived signature. Although originally derived from bulk transcriptomic data, this framework remains one of the few available approaches for transcriptomic subtyping of OCCC, in which the MesCC subtype has been associated with advanced-stage disease and poorer progression-free survival (27). Here, the behavior of the MesCC score suggested strongly that tissue context is important when bulk-derived signatures are applied to spatial data. MesCC was highest in stroma, intermediate in endometriosis and lowest in tumor. Although some inter-case heterogeneity was observed among tumor ROIs, no clear difference was evident according to ARID1A status. These findings may suggest that the MesCC signal partly reflects a broader mesenchymal or stromal context rather than simply tumor-intrinsic aggressiveness. Accordingly, the present findings do not necessarily contradict the MesCC framework but may indicate that its biological interpretation requires additional caution when applied to spatially resolved tissue regions.

The same caution applies to cell deconvolution analysis. The tumor microenvironment of OCCC is heterogeneous and spatially structured, and transcriptomic deconvolution may provide a useful overview of non-neoplastic cellular context. In this study, the estimated cell-type distribution was biologically plausible: stroma ROIs were fibroblast-rich, whereas endometriosis and tumor ROIs showed relatively stronger immune-associated signals. Tumor ROIs also showed relatively higher neutrophil and T-cell signals, including both CD4 and CD8, than endometriosis ROIs. The relatively clear compartmental differences among stromal, endometriotic and tumor regions were also broadly consistent with the metaprogram heatmap, suggesting that GeoMx-based spatial profiling may help resolve tissue-region-specific transcriptional features. However, these values remain model-based estimates derived from mixed regions and rather than precise cell-resolved measurements. Therefore, the present results are more appropriately interpreted as suggesting differences in surrounding cellular composition between compartments rather than providing a definitive description of the microenvironment.

Although this study represents a thorough analysis of carefully selected UK and Japanese cases in which endometriosis and ovarian clear cell carcinoma were adjacent on the same section, it has some potential shortcomings. The sample size was small, and cases were intentionally restricted to those in which tumor and adjacent endometriosis were present on the same section. This strengthened within-case spatial comparison but may limit generalizability. The analysis was performed on a region-of-interest basis because local tissue context can influence expression profiles in spatial transcriptomic analyses such as GeoMx. However, as the purpose of this study was to characterize spatially distinct transcriptional states in regions of tumor and endometriosis, rather than to classify whole cases, ROI-level analysis was considered most appropriate. The majority of the selected cases were FIGO stage I/II, so the conclusions are inevitably weighted toward early-stage disease. However, this should be interpreted in the context of the clinical characteristics of OCCC, which is often diagnosed at an early stage (6). Lastly, ARID1A classification was based on IHC rather than genomic characterization in all cases. However, previous studies have shown that ARID1A protein expression can be used to define *ARID1A* mutational status reliably (21). More generally, GeoMx provides very accurate spatial resolution and nucleus counting, which allow for derivation of proportions or relative abundance estimations, albeit not at single cell level. We nevertheless consider this a minor limitation in the context of this study, which aimed to assess major transcriptional differences between compartments and we showed that these data can be used also to assess tissue heterogeneity within metaprograms (29).

An important unresolved question is when, and in what way, these spatial transcriptional differences emerge during endometriosis-associated tumorigenesis and how they contribute to the malignant transformation from endometriosis to OCCC – the high overlap in gene expression within passage regions and immediately adjacent tumor and endometriosis suggests a graded transitional process. However, although *ARID1A* alterations are considered an early and important event in this process, our findings suggest that malignant transformation is unlikely to be explained by ARID1A loss alone, in keeping with mutation data (15).

Recent protein-based spatial profiling studies in OCCC suggest that tumor cell immune mimicry intersects with patterns of immune infiltration, with NK- and B-cell–like mimicry associated with improved prognosis in early-stage disease (30). Although paired profiling of adjacent endometriosis was not performed, these findings imply coordinated evolutionary changes in both tumor cells and the surrounding microenvironment, which may ultimately influence prognosis and therapeutic response. Together, they highlight a level of biological complexity that is unlikely to be captured by analyses focused on a limited set of genes.

Future studies should therefore integrate molecular events beyond ARID1A loss, including additional genetic, epigenetic and microenvironmental changes and the interplay among all of them to clarify the mechanisms underlying this transition. Further investigation of the relationship between the microenvironment and tumorigenesis may help determine whether immune, stromal and inflammatory features actively promote malignant transformation or instead reflect secondary changes associated with tumor development. Addressing these questions will require validation in larger, clinically annotated cohorts, as well as complementary approaches using single-cell and higher-resolution spatial technologies and *in vivo* models. Such studies will also be important for assessing the prognostic and therapeutic relevance of the spatially defined biological differences identified in the present study and how the immune microenvironment might be used as a therapeutic target in these tumors. In fact, although immune checkpoint inhibitors (ICI) have shown some evidence of clinical activity, for example in the PEACOCC trial (31), their efficacy as monotherapy in ovarian clear cell carcinoma remains modest, and robust predictive biomarkers are still lacking. In contrast, combination strategies involving ICI (32) or targeting other immune cells such as neutrophils might be an interesting therapeutic approach in this type of tumors and a more granular immune-based stratification may enable the identification of prognostically and predictively distinct subgroups (30,33).

In summary, we identify pronounced transcriptomic differences between tumor tissue and adjacent endometriosis, suggesting progressive molecular evolution during malignant transformation. This compartmental difference was consistently observed across differential expression, pathway, immune context and meta-program analyses, and was independent of geographic origin or ARID1A status. Collectively, these findings suggest that OCCC establishes transcriptional programs that are distinct from adjacent endometriosis, reflecting concomitant alterations in the surrounding microenvironment.

## Acknowledgements

This work was funded by Ovarian Cancer Action, Cancer Research UK and Imperial NIHR Biomedical Research Centre. We acknowledge support from the Imperial College Healthcare Tissue Bank and the Imperial BRC Genomics Facility. The spatial transcriptomics of Japanese samples were performed by Eurofins GeneticLab, Sapporo, Japan. GG was supported by an ESMO Translational Fellowship and an AIRC Gianni Bonadonna Fellowship. IMcN is an NIHR Senior Investigator.

## Supplementary Figures Legends

**Figure S1. Normalization**. Assessment of normalization in UK (left) and Japanese (right) cohorts. **A**. Relationship between Q3 values and negative probe geometric mean counts across ROIs. Each point represents one ROI and is colored by tissue category. The dashed line indicates the identity line. **B**. boxplots showing raw counts **C**, normalized counts across segments.

**Figure S2. UMAP views of ROIs by cohort**. UMAP embeddings showing ROI distributions in the UK (left) and Japan (right) cohorts before combat batch correction. Points represent individual ROIs and are colored by tissue category.

**Figure S3. Assessment of cohort-associated effects before and after batch correction**. Linear regression analysis of housekeeping gene expression comparing UK and Japan cohorts. Triangles indicate values before correction and circles after correction.

**Figure S4. KEGG pathway enrichment for tumor versus endometriosis DEGs**. Dot plots showing KEGG pathway enrichment of tumor-upregulated genes (left) and endometriosis-upregulated genes (right) in the combined, UK and Japan analyses. Dot color indicates adjusted *P*-value and dot size indicates gene ratio.

**Figure S5. Comparison between tumor and endometriosis using the original cohort datasets prior to batch correction**. Volcano plots and pathway enrichment analyses for tumor versus endometriosis. The upper panels show differential gene expression, the middle panels show representative GO Biological Process terms, and the lower panels show KEGG pathway enrichment results. The left panels represent the UK cohort, and the right panels represent the Japanese cohort.

**Figure S6. Passage-associated comparisons after batch correction**. Volcano plots showing differential expression for tumor versus tumor-passage (left) and endometriosis versus endometriosis-passage (right) ROIs.

**Figure S7. Comparisons of endometriosis and tumor against stroma**. Volcano plots and pathway enrichment analyses for tumor versus stroma (left) and endometriosis versus stroma (right). **A** differential expression; **B**. representative GO Biological Process; **C**. KEGG enrichment results.

**Figure S8. Meta-program scores across ROI groups**. Heatmap showing mean meta-program scores across ROI groups. Rows represent individual meta-programs and columns represent ROI groups.

**Figure S9. ARID1A transcription in tumor ROIs**. *ARID1A* transcription in ARID1A-retained and ARID1A-loss tumor ROIs (left) and in tumor ROIs from the UK and Japan cohorts (right).

**Figure S10. ARID1A-stratified KEGG pathway enrichment**. Dot plots showing KEGG pathway enrichment of tumor-upregulated genes (left) and endometriosis-upregulated genes (right) in the combined analysis and in ARID1A-loss and ARID1A-retained groups. Dot color indicates adjusted *P*-value and dot size indicates gene ratio.

**Figure S11. Case-level and stratified MesCC score analyses**. MesCC score summaries across individual cases. Points represent individual ROIs and box plots summarize group distributions.

Statistical comparisons were performed using the Kruskal–Wallis test for the case-level analysis.

**Figure S12. ROI-level SpatialDecon profiles by compartment**. Stacked bar plots showing SpatialDecon-estimated relative non-tumor cell-type proportions in individual tumor (left) endometriosis (middle) and stroma (right) ROIs.

**Figure S13. Cohort-stratified and ARID1A-stratified SpatialDecon summaries**. SpatialDecon-estimated relative non-tumor cell-type proportions across compartments, stratified by cohort (left) and ARID1A status (right).

## References

1. Glasspool RM, McNeish IA. Clear Cell Carcinoma of Ovary and Uterus. Current oncology reports 2013;16(6):566–72 doi 10.1007/s11912-013-0346-0.

2. Rosso R, Turinetto M, Borella F, Chopin N, Meeus P, Lainè A, et al. Ovarian clear cell carcinoma: open questions on the management and treatment algorithm. The oncologist 2025;30(1) doi 10.1093/oncolo/oyae325.

3. Okamoto A, Glasspool RM, Mabuchi S, Matsumura N, Nomura H, Itamochi H, et al. Gynecologic Cancer InterGroup (GCIG) Consensus Review for Clear Cell Carcinoma of the Ovary. Int J Gynecol Cancer 2014;24(9 Suppl 3):S20–s5 doi 10.1097/igc.0000000000000289.

4. Wei YF, Ning L, Xu YL, Ma J, Li DR, Feng ZF, et al. Worldwide patterns and trends in ovarian cancer incidence by histological subtype: a population-based analysis from 1988 to 2017. EClinicalMedicine 2025;79:102983 doi 10.1016/j.eclinm.2024.102983.

5. Kuroda T, Kohno T. Precision medicine for ovarian clear cell carcinoma based on gene alterations. International journal of clinical oncology/Japan Society of Clinical Oncology 2020;25(3):419–24 doi 10.1007/s10147-020-01622-z.

6. Iida Y, Okamoto A, Hollis RL, Gourley C, Herrington CS. Clear cell carcinoma of the ovary: a clinical and molecular perspective. Int J Gynecol Cancer 2021;31(4):605–16 doi 10.1136/ijgc-2020-001656.

7. Suda K, Nakaoka H, Yoshihara K, Ishiguro T, Tamura R, Mori Y, et al. Clonal Expansion and Diversification of Cancer-Associated Mutations in Endometriosis and Normal Endometrium. Cell reports 2018;24(7):1777–89 doi 10.1016/j.celrep.2018.07.037.

8. Shigetomi H, Tsunemi T, Haruta S, Kajihara H, Yoshizawa Y, Tanase Y, et al. Molecular mechanisms linking endometriosis under oxidative stress with ovarian tumorigenesis and therapeutic modalities. Cancer investigation 2012;30(6):473–80 doi 10.3109/07357907.2012.681821.

9. Pearce CL, Templeman C, Rossing MA, Lee A, Near AM, Webb PM, et al. Association between endometriosis and risk of histological subtypes of ovarian cancer: a pooled analysis of case-control studies. Lancet Oncol 2012;13(4):385–94 doi S1470-2045(11)70404-1[pii] 10.1016/S1470-2045(11)70404-1 [doi].

10. Jiang W, Xu W, Chen F. Dysfunction of natural killer cells promotes immune escape and disease progression in endometriosis. Frontiers in immunology 2025;16:1657605 doi 10.3389/fimmu.2025.1657605.

11. Shifon S, Tyrinova T, Veretelnikova T, Pasman N, Chernykh E. Endometriosis as an immune-mediated disease: pathogenetic mechanisms and therapeutic strategies. Frontiers in immunology 2025;16:1727183 doi 10.3389/fimmu.2025.1727183.

12. Jones S, Wang TL, Shih Ie M, Mao TL, Nakayama K, Roden R, et al. Frequent mutations of chromatin remodeling gene ARID1A in ovarian clear cell carcinoma. Science 2010;330(6001):228–31 doi 10.1126/science.1196333.

13. Wiegand KC, Shah SP, Al-Agha OM, Zhao Y, Tse K, Zeng T, et al. ARID1A mutations in endometriosis-associated ovarian carcinomas. N Engl J Med 2010;363(16):1532–43 doi 10.1056/NEJMoa1008433.

14. Anglesio MS, Papadopoulos N, Ayhan A, Nazeran TM, Noe M, Horlings HM, et al. Cancer-Associated Mutations in Endometriosis without Cancer. N Engl J Med 2017;376(19):1835–48 doi 10.1056/NEJMoa1614814.

15. Anglesio MS, Bashashati A, Wang YK, Senz J, Ha G, Yang W, et al. Multifocal endometriotic lesions associated with cancer are clonal and carry a high mutation burden. J Pathol 2015;236(2):201–9 doi 10.1002/path.4516.

16. Maeda D, Mao TL, Fukayama M, Nakagawa S, Yano T, Taketani Y, et al. Clinicopathological significance of loss of ARID1A immunoreactivity in ovarian clear cell carcinoma. International journal of molecular sciences 2010;11(12):5120–8 doi 10.3390/ijms11125120.

17. Luchini C, Veronese N, Solmi M, Cho H, Kim JH, Chou A, et al. Prognostic role and implications of mutation status of tumor suppressor gene ARID1A in cancer: a systematic review and meta-analysis. Oncotarget 2015;6(36):39088–97 doi 10.18632/oncotarget.5142.

18. Iida Y, Churchman M, Hollis RL, Taylor S, Bartos C, Croy I, et al. Combined genomic and molecular analysis defines prognostic markers of relapse in stage IA-IC1 clear cell ovarian carcinoma. Gynecol Oncol 2025;198:66–74 doi 10.1016/j.ygyno.2025.05.016.

19. Fonseca MAS, Haro M, Wright KN, Lin X, Abbasi F, Sun J, et al. Single-cell transcriptomic analysis of endometriosis. Nat Genet 2023;55(2):255–67 doi 10.1038/s41588-022-01254-1.

20. Guo Q, Chen X, Bi R, Li H, Liu M, Ju X, et al. Epithelial characteristics of ovarian clear cell carcinoma at single-cell resolution. Commun Biol 2025;8(1):1156 doi 10.1038/s42003-025-08617-4.

21. Khalique S, Naidoo K, Attygalle AD, Kriplani D, Daley F, Lowe A, et al. Optimised ARID1A immunohistochemistry is an accurate predictor of ARID1A mutational status in gynaecological cancers. The journal of pathology Clinical research 2018;4(3):154–66 doi 10.1002/cjp2.103.

22. Kawabata A, Yanaihara N, Nagata C, Saito M, Noguchi D, Takenaka M, et al. Prognostic impact of interleukin-6 expression in stage I ovarian clear cell carcinoma. Gynecol Oncol 2017;146(3):609–14 doi 10.1016/j.ygyno.2017.06.027.

23. Zhang Y, Parmigiani G, Johnson WE. ComBat-seq: batch effect adjustment for RNA-seq count data. NAR Genom Bioinform 2020;2(3):lqaa078 doi 10.1093/nargab/lqaa078.

24. Ma H, Srivastava S, Ho SWT, Xu C, Lian BSX, Ong X, et al. Spatially Resolved Tumor Ecosystems and Cell States in Gastric Adenocarcinoma Progression and Evolution. Cancer Discov 2025;15(4):767–92 doi 10.1158/2159-8290.Cd-24-0605.

25. Gavish A, Tyler M, Greenwald AC, Hoefflin R, Simkin D, Tschernichovsky R, et al. Hallmarks of transcriptional intratumour heterogeneity across a thousand tumours. Nature 2023;618(7965):598–606 doi 10.1038/s41586-023-06130-4.

26. Hwang I, Cho Y, Kang SY, Kim DG, Ahn S, Lee J, et al. Comparative analysis of ARID1A mutations with mRNA levels and protein expression in gastric carcinoma. Pathology, research and practice 2024;255:155063 doi 10.1016/j.prp.2023.155063.

27. Tan TZ, Ye J, Yee CV, Lim D, Ngoi NYL, Tan DSP, et al. Analysis of gene expression signatures identifies prognostic and functionally distinct ovarian clear cell carcinoma subtypes. EBioMedicine 2019;50:203–10 doi 10.1016/j.ebiom.2019.11.017.

28. Danaher P, Kim Y, Nelson B, Griswold M, Yang Z, Piazza E, et al. Advances in mixed cell deconvolution enable quantification of cell types in spatial transcriptomic data. Nat Commun 2022;13(1):385 doi 10.1038/s41467-022-28020-5.

29. Gaspard-Boulinc LC, Gortana L, Walter T, Barillot E, Cavalli FMG. Cell-type deconvolution methods for spatial transcriptomics. Nature reviews Genetics 2025;26(12):828–46 doi 10.1038/s41576-025-00845-y.

30. Tai YT, Lin WC, Ye J, Chen DT, Chen KC, Wang DY, et al. Spatial Profiling of Ovarian Clear Cell Carcinoma Reveals Immune-Hot Features. Mod Pathol 2025;38(1):100630 doi 10.1016/j.modpat.2024.100630.

31. Kristeleit R, Devlin MJ, Clamp A, Gourley C, Roux R, Hall M, et al. Pembrolizumab in Patients With Advanced Clear Cell Gynecological Cancer: A Phase 2 Nonrandomized Clinical Trial. JAMA Oncol 2025;11(4):377–85 doi 10.1001/jamaoncol.2024.6797.

32. Peng Z, Li H, Gao Y, Sun L, Jiang J, Xia B, et al. Sintilimab combined with bevacizumab in relapsed or persistent ovarian clear cell carcinoma (INOVA): a multicentre, single-arm, phase 2 trial. Lancet Oncol 2024;25(10):1288–97 doi 10.1016/s1470-2045(24)00437-6.

33. Koenderman L, Vrisekoop N. Neutrophils in cancer: from biology to therapy. Cell Mol Immunol 2025;22(1):4–23 doi 10.1038/s41423-024-01244-9.

