## Supplementary methods and figures for "Spatial transcriptomic analysis reveals coordinated gene expression in ovarian clear cell carcinoma and adjacent endometriosis in UK and Japanese patients"

##### GeoMx transcriptomic analysis and quality control

GeoMx transcriptomic profiling was performed using the Human NGS Whole Transcriptome Atlas RNA panel. Spatial transcriptomic analysis was conducted independently for the UK and JP cohorts. A total of 136 ROIs from the UK cohort and 144 ROIs from the JP cohort were analyzed.

The data were processed in R using a workflow based on Biobase and GeoMx-related packages. Segment-level quality control thresholds were applied as follows: minimum segment reads, 1,000; percent trimmed, 80; percent stitched, 80; percent aligned, 80; sequencing saturation, 50; minimum negative count, 1; maximum NTC count, 9,000; minimum nuclei, 20; and minimum area, 1,000. Probe-level quality control was then performed using a minimum probe ratio of 0.1 and percentFailGrubbs of 20, followed by local outlier removal and aggregation of probe-level information to the target level. These QC parameters were applied to both the UK and JP cohorts.

The limit of quantification (LOQ) was calculated based on negative control probes, using a cutoff parameter of 2 and a minimum LOQ of 2. For each ROI, the number of genes exceeding the LOQ was calculated and a segment-level gene detection rate was derived. ROIs with a gene detection rate of at least 0.05 were retained for downstream analysis. At the gene level, endogenous genes were retained if they were detected above the LOQ in at least 1% of ROIs within each cohort after QC. Negative control probes were used for LOQ calculation, QC assessment and evaluation of normalization methods.

Expression data were normalized using 75th percentile normalization (Q3 normalization). To assess the appropriateness of Q3 normalization, negative-probe normalization was also calculated and expression distributions and between-ROI variability were compared. Based on these results, Q3-normalized values were used for downstream analyses. For visualization and dimensionality reduction analyses, expression values were log2-transformed as log2 (Q3 + 1).

##### Cohort integration and batch correction

Before integrated analysis of the UK and Japanese cohorts, only genes commonly measured in both cohorts were retained. Q3-normalized expression values from each cohort were log2-transformed [log2(Q3 + 1)], after which a single expression matrix consisting of shared genes was generated and the corresponding metadata were merged. The structure of the integrated data before correction was assessed using UMAP. Batch correction was then performed using ComBat, with cohort (UK/JP) specified as the batch variable and UMAP was rerun under the same conditions to assess changes in ROI distribution before and after correction.

Residual batch effects were evaluated using stroma ROIs. First, the top 20 most variable genes among stroma ROIs were selected and principal component analysis (PCA) was performed before and after correction using these genes together with the subset present in the data from 10 housekeeping genes (*ACTB, GAPDH, RPL19, RPL37A, RPLP0, TUBB, CLTC, GUSB, HPRT1 and PPIA*). In addition, gene-wise linear regression models were fitted to housekeeping gene expression in stroma ROIs with cohort as the explanatory variable and effect sizes (β) and P-values for JP relative to UK were estimated. These analyses were performed both before and after ComBat correction and P-values were adjusted for multiple testing using the Benjamini–Hochberg method.

**Normalization**

Q3 counts showed a positive relationship with negative probe GeoMean counts in both cohorts (Figure S1A). Raw count distributions varied across segments, with noticeable differences in overall signal intensity and spread. After Q3 normalization, the distributions became more closely aligned across segments, with more comparable medians and interquartile ranges. These findings indicate that Q3 normalization reduced segment-to-segment technical variability (Figure S1B and C). These QC metrics supported downstream analyses using batch-corrected expression values in the combined dataset.

**Meta-program Scoring**

For heatmap visualization, the mean score of each MP was first calculated for each ROI group. This group-averaged MP score matrix was then further row-wise Z-score standardized so that relative differences among ROI groups could be visualized for each MP. In the main analysis, Stroma, Endometriosis and Tumor were displayed and evaluated. Accordingly, the heatmap color scale represents relative patterns across groups for each MP rather than raw expression intensity.

Figure S1

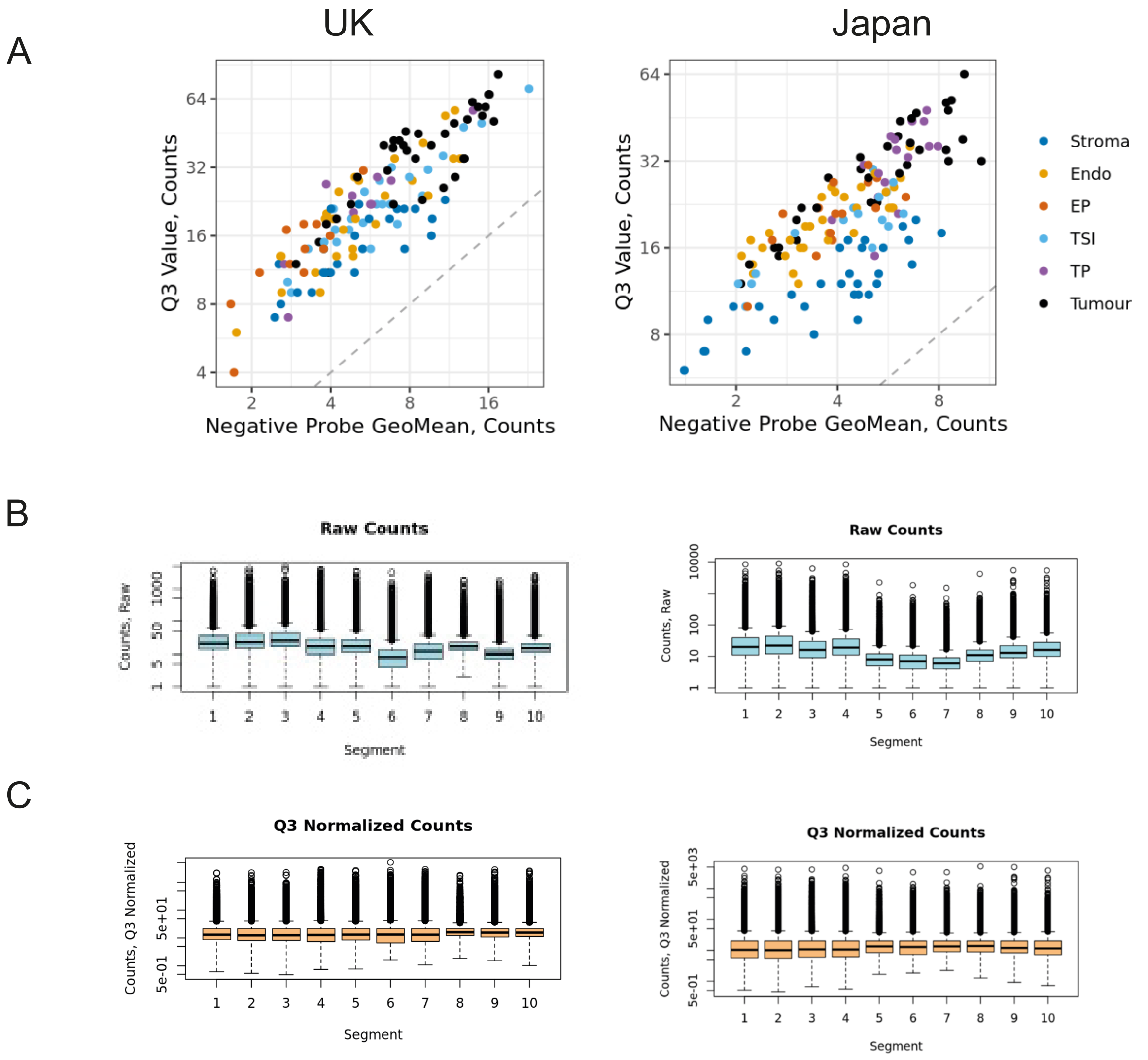


Figure S2

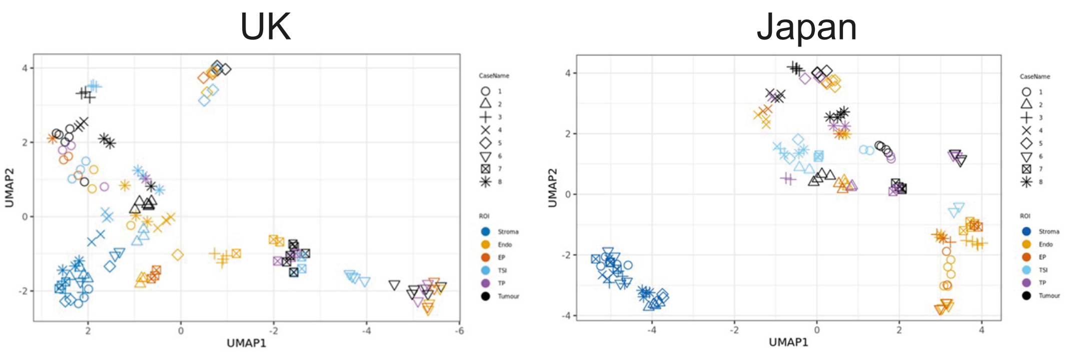


Figure S3

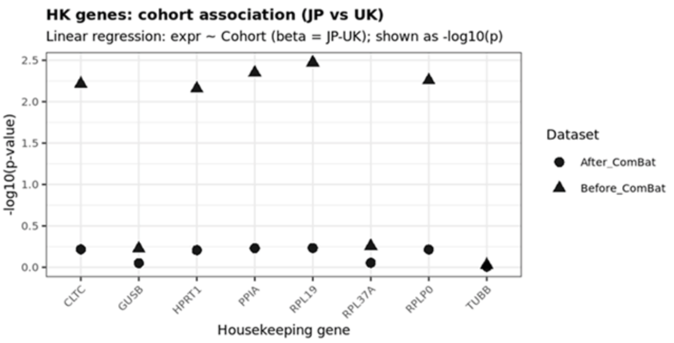

### Figure S4

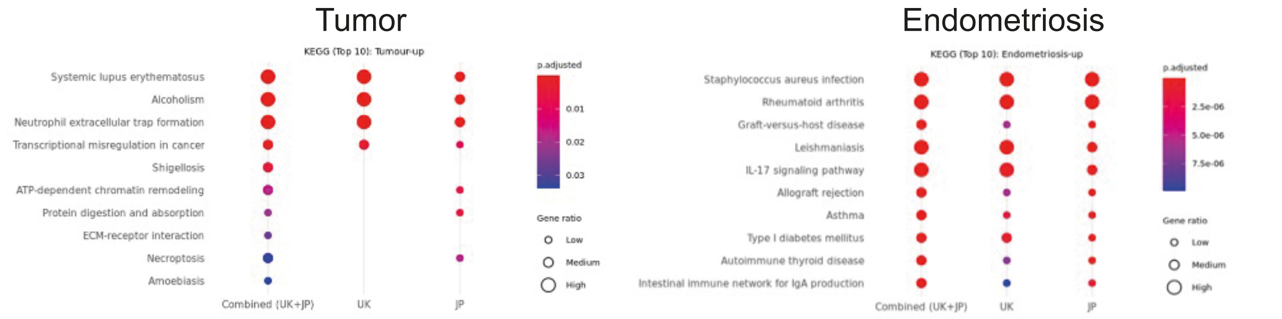


### Figure S5

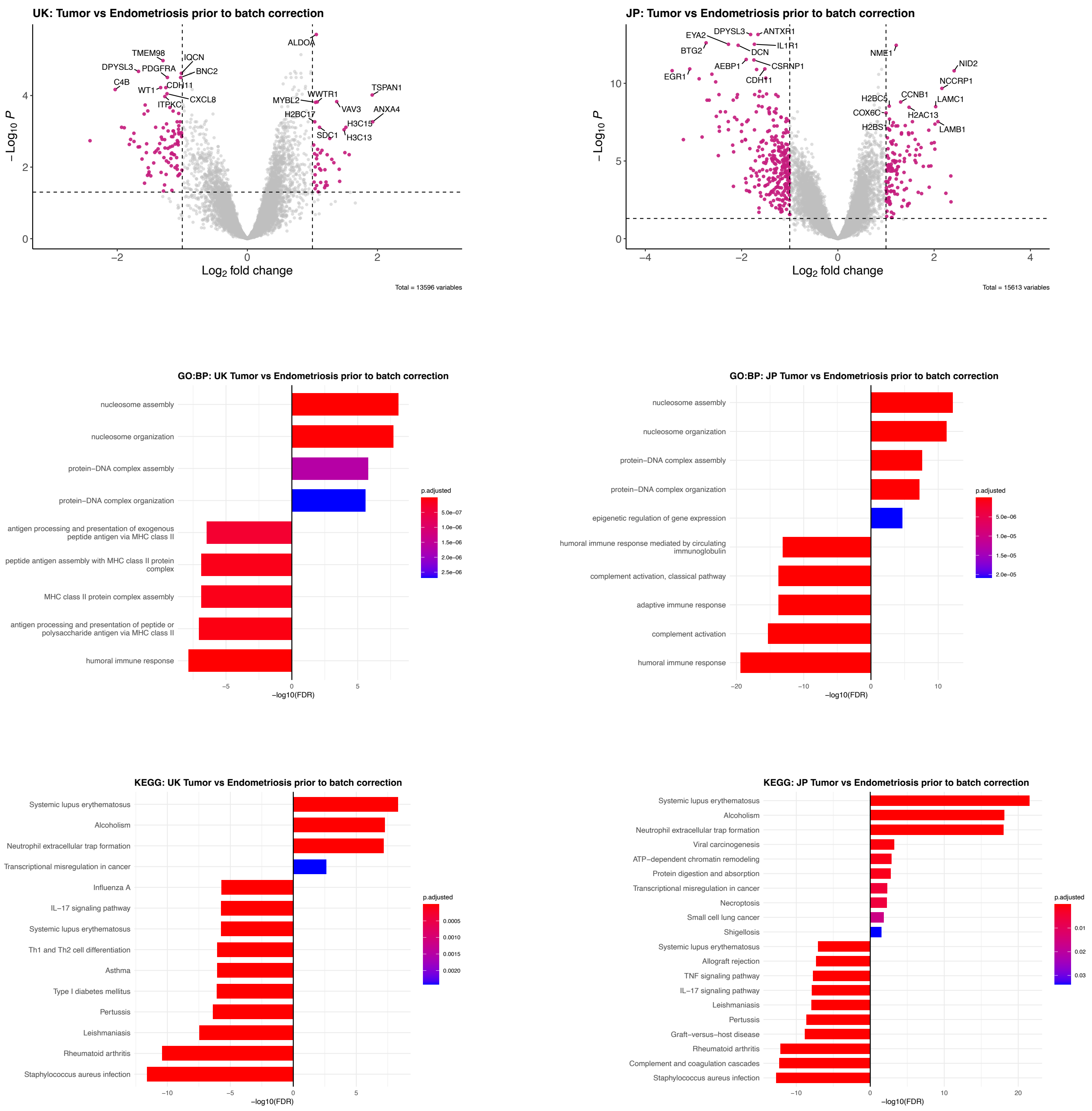

#### Figure S6

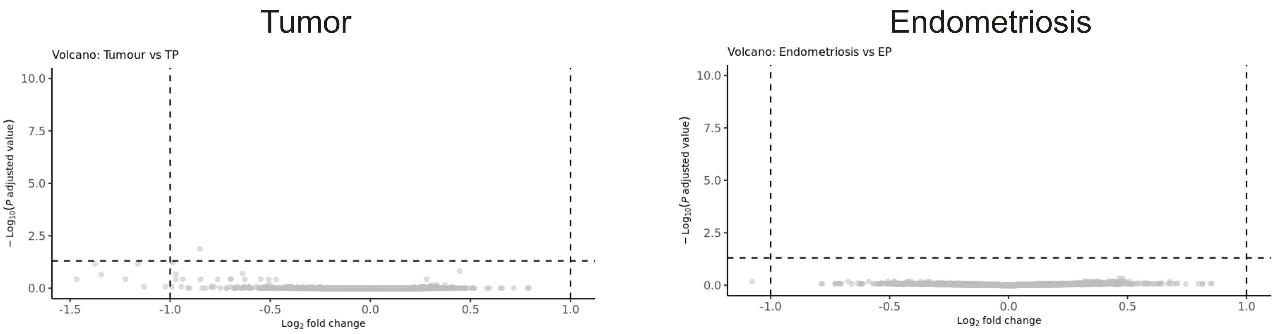


#### Figure S7

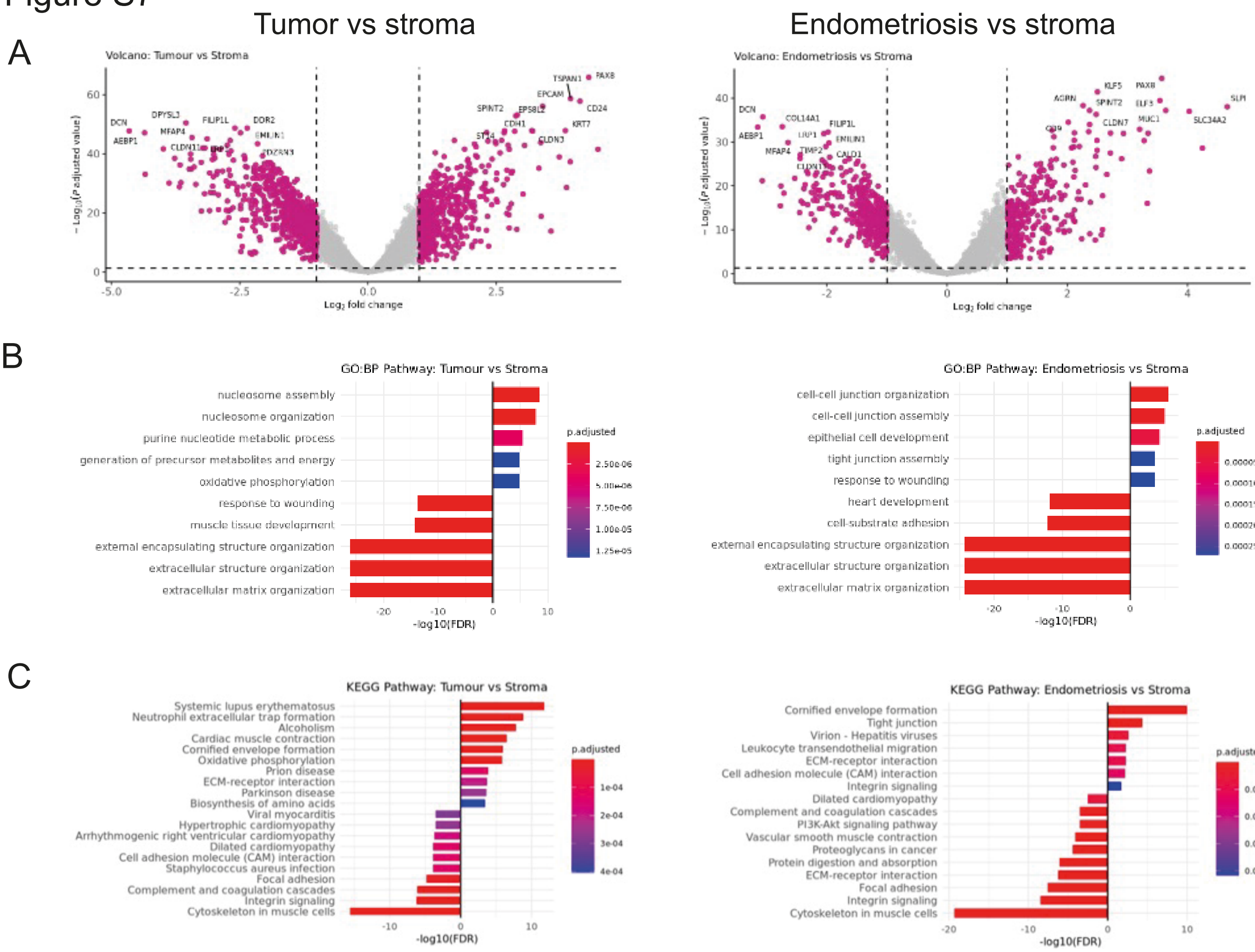


#### Figure S8

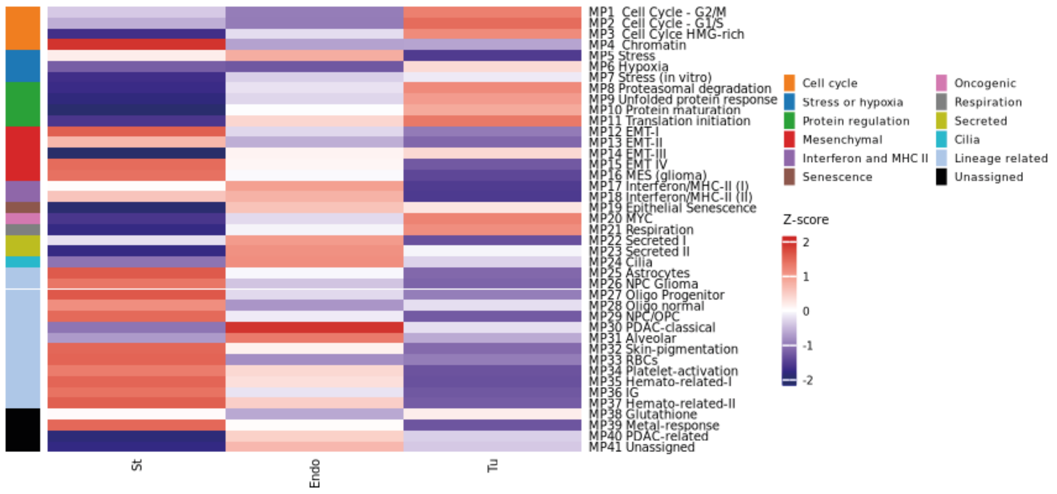

### Figure S9

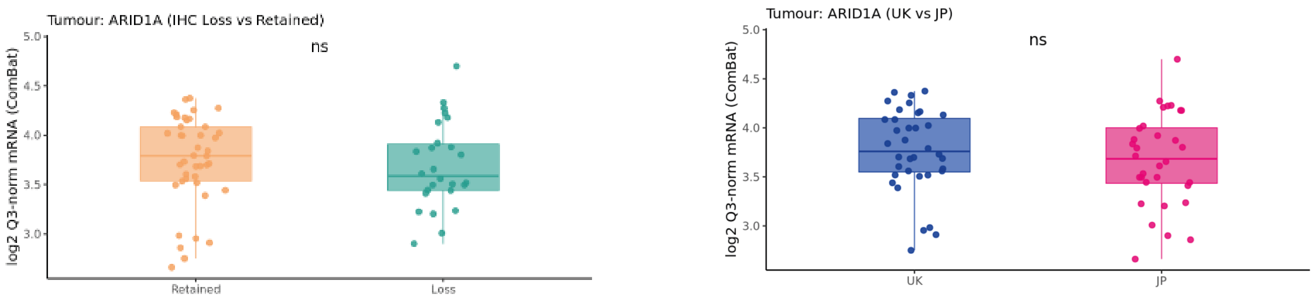


### Figure S10

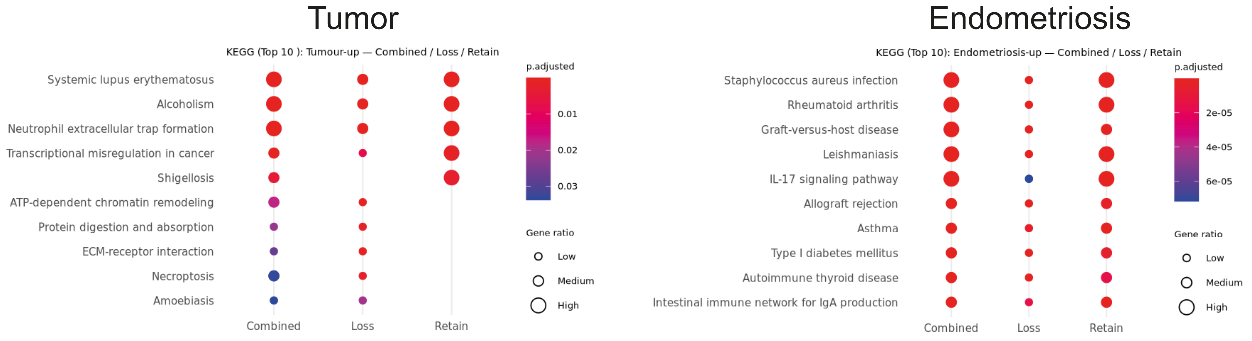


### Figure S11

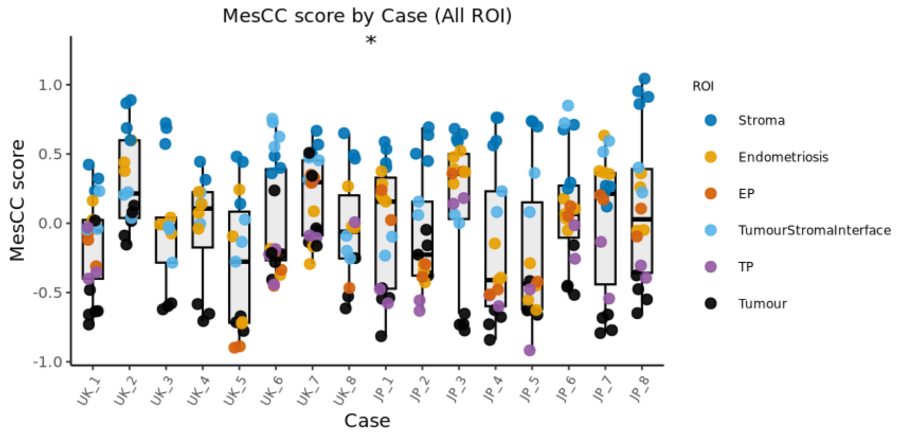


### Figure S12

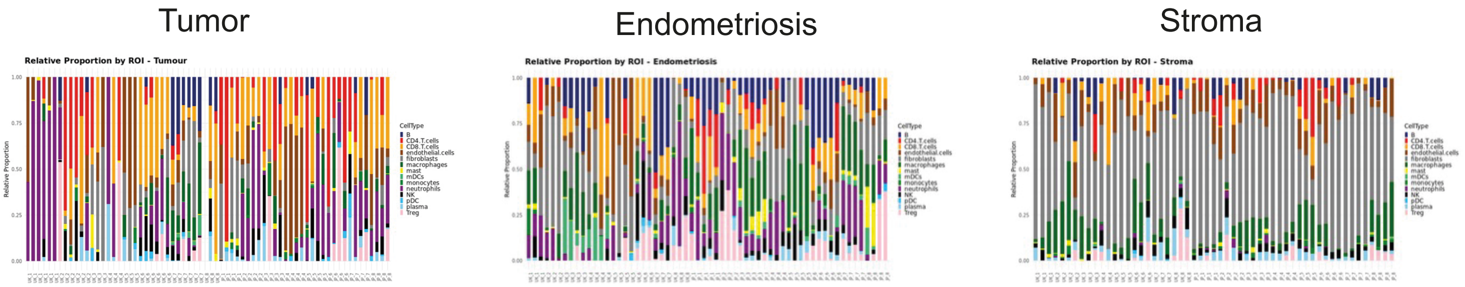

### Figure S13

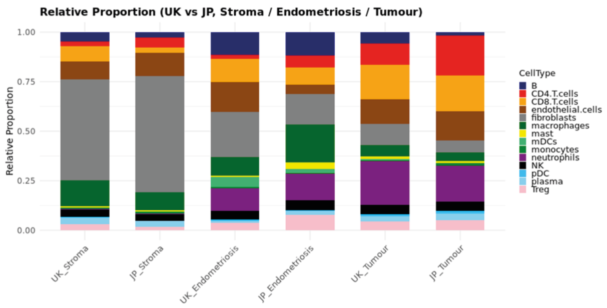
Relative Proportion (UK vs JP, Stroma / Endometriosis / Tumour)
Relative Proportion
1.00
0.75
0.50
0.25
0.00
UK_Stroma
JP_Stroma
UK_Endometriosis
JP_Endometriosis
UK_Tumour
JP_Tumour
CellType
B
CD4.T.cells
CD8.T.cells
endothelial.cells
fibroblasts
macrophages
mast
mDCs
monocytes
neutrophils
NK
pDC
plasma
Treg

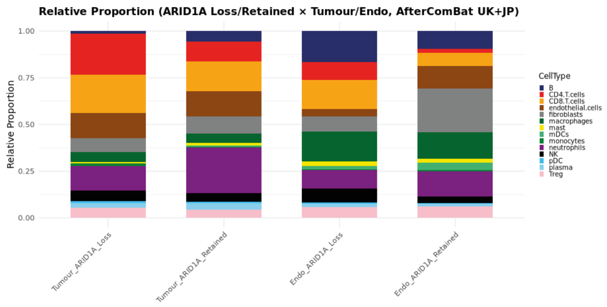
Relative Proportion (ARID1A Loss/Retained × Tumour/Endo, AfterComBat UK+JP)
Relative Proportion
1.00
0.75
0.50
0.25
0.00
Tumour_ARID1A_Loss
Tumour_ARID1A_Retained
Endo_ARID1A_Loss
Endo_ARID1A_Retained
CellType
B
CD4.T.cells
CD8.T.cells
endothelial.cells
fibroblasts
macrophages
mast
mDCs
monocytes
neutrophils
NK
pDC
plasma
Treg
